# SARS-CoV-2 infects brain choroid plexus and disrupts the blood-CSF-barrier

**DOI:** 10.1101/2020.08.20.259937

**Authors:** Laura Pellegrini, Anna Albecka, Donna L. Mallery, Max J. Kellner, David Paul, Andrew P. Carter, Leo C. James, Madeline A. Lancaster

## Abstract

Coronavirus disease-19 (COVID-19), caused by the SARS-CoV-2 virus, leads primarily to respiratory symptoms that can be fatal, particularly in at risk individuals. However, neurological symptoms have also been observed in patients, including headache, seizures, stroke, and fatigue. The cause of these complications is not yet known, and whether they are due to a direct infection of neural cells, such as neurons and astrocytes, or through indirect effects on supportive brain cells, is unknown. Here, we use brain organoids to examine SARS-CoV-2 neurotropism. We examine expression of the key viral receptor ACE2 in single-cell RNA sequencing (scRNA-seq) revealing that only a subset of choroid plexus cells but not neurons or neural progenitors express this entry factor. We then challenge organoids with both SARS-CoV-2 spike protein pseudovirus and live virus to demonstrate high viral tropism for choroid plexus epithelial cells but not stromal cells, and little to no infection of neurons or glia. We find that infected cells of the choroid plexus are an apolipoprotein and ACE2 expressing subset of epithelial barrier cells. Finally, we show that infection with live SARS-CoV-2 leads to barrier breakdown of the choroid plexus. These findings suggest that neurological complications may result from effects on the choroid plexus, an important barrier that normally prevents entry of immune cells and cytokines into the cerebrospinal fluid (CSF) and brain.

## Introduction

The COVID-19 global pandemic caused by the Severe Acute Respiratory Syndrome Coronavirus 2 (SARS-CoV-2) has infected more than 20 million people and caused around 740,000 deaths as of early August 2020 (worldometers.info/coronavirus) and it still poses a constant threat to public health systems worldwide. Respiratory symptoms, predominantly associated with the infection, include fever, chest tightness and persistent cough (Wu et al., 2020). SARS-CoV-2 shares around 80% sequence similarity with SARS-CoV (Wang et al., 2020) and both use their spike (S) glycoprotein to mediate host cell entry (Wang et al., 2020; Wu et al., 2020).

SARS-CoV-2 shares around 80% sequence similarity with SARS-CoV (Wang et al., 2020) and both use their spike (S) glycoprotein to mediate host cell entry (Wang et al., 2020; Wu et al., 2020). SARS-CoV-2 enters the cells using a cell surface receptor, angiotensin-converting enzyme 2 (ACE2) (Hoffmann et al., 2020b, 2020a). Co-entry factors such as TMPRSS2 (Hoffmann et al., 2020a), TMPRSS4 (Zang et al., 2020) and neuropilin1 (Cantuti-Castelvetri et al., 2020; Daly et al., 2020) have also been reported to potentiate infectivity. CoVs are widely believed to enter the cells via two mechanisms: via the endocytic pathway and via a non-endosomal pathway (Ou et al., 2020; Yang and Shen, 2020). Direct fusion of the virus to the plasma membrane seems to be the predominant route of entry of SARS-CoV-2 (Hoffmann et al., 2020b). SARS-CoV-2 and SARS-CoV both use their S protein, composed of two subunits S1 and S2, for viral entry: S1 facilitates viral attachment to the receptor ACE2 and S2 is essential for membrane fusion, which also involves the host cell protease TMPRSS2 (Hoffmann et al., 2020b, 2020a; Walls et al., 2020; Wrapp et al., 2020).

The overwhelming majority of respiratory symptoms suggest that the first target for viral entry is the respiratory tract (Sungnak et al., 2020). Recent RNA-sequencing data from multiple tissues from healthy human donors reported viral tropism for nasal epithelial cells, predominantly ciliated cells and goblet cells, highly secretory cells involved in mucus secretion, expressing viral entry-associated genes including ACE2 and TMPRSS2 (Sungnak et al., 2020). Expression of these viral entry genes has also been reported to promote viral entry in human small intestine enterocytes (Zang et al., 2020), suggesting potential viral spread through the fecal-oral route and explaining the common gastrointestinal symptoms observed in patients. These findings have been corroborated by experiments conducted in human intestinal organoids, which show high expression of SARS-CoV-2 entry factors and infection of human enterocytes with both SARS-CoV and SARS-CoV-2 (Lamers et al., 2020). Enterocytes are highly secretory cells which express ACE2 and produce large lipid vesicles containing lipoproteins involved in the generation of chylomicrons, trygliceride-rich lipoprotein particles important for systemic lipid transfer (Ko et al., 2020).

Even though respiratory systems seem to be the most affected, increasing evidence from clinical reports indicates a rise in both acute and chronic neurological symptoms including sudden and complete loss of smell, headache, seizures, stroke and long-term clinical complications such as persistent fatigue (Montalvan et al., 2020) and meningitis/encephalitis (Moriguchi et al., 2020). These symptoms suggest viral tropism for brain cells; however, which cell types remains to be fully determined. Furthermore, while preliminary reports using in vitro systems, including neural organoids, suggest some neurotropism, the physiological relevance is still unclear. In particular, the degree of infection relative to more susceptible cell types, as well as the route of entry in to the brain, remains to be explored.

Interestingly, one case reported SARS-CoV-2 in the cerebrospinal fluid (CSF) but not in the nasopharyngeal specimen collected from a patient presenting neurological symptoms (Moriguchi et al., 2020). Dissemination through the central nervous system (CNS), however, has not yet been demonstrated, and viral presence in brain tissue and CSF has not been widely reported. Nonetheless, viral entry through the olfactory bulb and cribriform plate have been suggested as potential routes of entry to the CNS (Montalvan et al., 2020). Other possible routes of entry of virus or bacteria into the brain are through the blood-CNS barriers such as the blood-brain-barrier (BBB) or the blood-CSF-barrier (B-CSF-B). The BBB, which separates the systemic blood from the brain parenchyma, is a complex barrier constituted by multiple cells types and mainly formed by the tight junctions between endothelial cells. The B-CSF-B is formed by the epithelial cells of the choroid plexus (ChP) and separates the blood from the cerebrospinal fluid (CSF) (Ghersi-Egea et al., 2018; Lehtinen et al., 2011; Lun et al., 2015a; Strazielle and Ghersi-Egea, 2013). The ChP also produces the CSF, a colourless fluid that provides essential nutrients and helps clear out waste byproducts (Dani et al., 2019; Fame and Lehtinen, 2020; Ghersi-Egea et al., 2018; Lehtinen and Walsh, 2011; Lehtinen et al., 2011; Lun et al., 2015b). CSF is produced by filtering blood from a dense network of fenestrated capillaries that lie within the stroma. The stroma is a rich environment that also provides a site of immune surveillance. But it is the epithelial cell layer of the ChP that forms the tight barrier and prevents entry of pathogens and toxic substances into the CSF (Lun et al., 2015a), as well as acting as a gateway for immune cells (Schwerk et al., 2015).

Because human brain tissue is difficult to access for the purpose of functional studies, including the investigation of viral tropism, recent studies have made use of 3D in vitro models called cerebral organoids to examine infection, for example with Zika virus (Qian et al., 2017; Zhou et al., 2017). These tissues can faithfully recapitulate, on a cellular-architecture and on a gene expression level, the early stages of brain development and can form mature neurons organised in thick axon tracts when cultured at the air-liquid interface (Giandomenico et al., 2019; Lancaster et al., 2017). We recently developed a cerebral organoid model to study the ChP (Pellegrini et al., 2020), which recapitulates the epithelial polarisation of ChP cells and the formation of a tight barrier which separates the surrounding media from the CSF-like fluid secreted by the ChP (Lancaster et al., 2017, 2013). To test viral tropism of SARS-CoV-2 in various cells of the CNS, we examined the expression patterns of viral entry factors in cerebral and ChP organoids, and tested for infection with both pseudovirions carrying SARS-CoV-2 spike, and live SARS-CoV-2. We found that ChP expressed SARS-CoV-2 entry factors in a previously unidentified lipoprotein-producing population of cells and that SARS-CoV-2 spike pseudovirions and live virus can productively infect ChP epithelial cells. In contrast, neurons and other CNS cell types were not generally susceptible, except under extreme infection with large viral quantities. We observed that the primary effect of the virus was on ChP cells, which disrupted integrity of this key CNS barrier.

## Results

### ACE2 and other entry factors are expressed in the choroid plexus

To assess whether SARS-CoV-2 entry factors are present in various cell types in organoids, we looked at the expression of the receptor ACE2 and the co-entry factor TMPRSS2 in different clusters of cells from previously published scRNA-seq data from ChP and telencephalic organoids (Pellegrini et al., 2020) (Fig. 1A, Fig. S1A). Expression of ACE2 and TMPRSS2 was detected predominantly in the immature ChP/hem cluster, and mature ChP clusters, but not in the neural progenitor or neuron clusters. ACE2 was also detected, although to a lesser extent in the ChP stroma (Fig. 1A, Fig. S1A). To examine whether these results are in agreement with the expression in vivo, we analysed data from the Allen Brain Atlas (Hawrylycz et al., 2012; Miller et al., 2017) reporting expression levels of ACE2 in different human brain regions (Fig. 1B). Among all the different brain regions compared, we found highest levels of ACE2 in the ChP (Fig. 1B), validating our findings in vitro.

**Figure 1.**
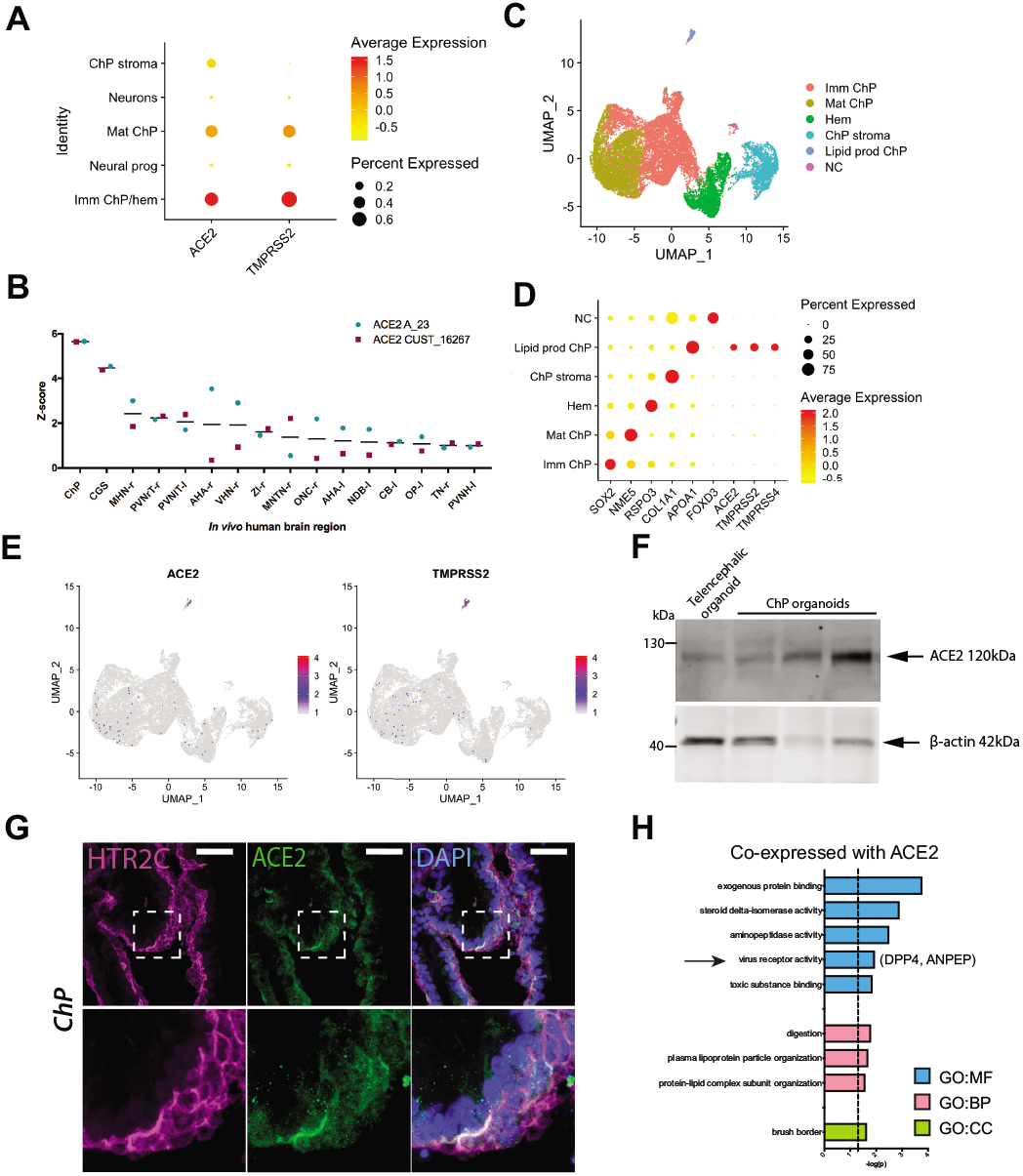
ACE2 and other entry factors are expressed in the choroid plexus. (A) Dot plot showing average expression and percentage of cells expressing SARS-CoV-2 entry factors ACE2 and TMRPSS2 in the five main clusters identified by scRNA-seq of ChP and telencephalic organoids (Pellegrini et al., 2020). (B) Allen Brain Atlas expression data of ACE2 (probe names: ACE2 A_23 and ACE2 CUST_16267) in different human brain regions, with highest expression in the ChP. Regions with an average z-score across the two probes (black line) of greater than 1 are shown. (C) UMAP plot showing subclustering of all ChP cell types identified by scRNA-seq. Imm ChP = immature ChP; Mat ChP = mature ChP, Lipid prod ChP = lipid producing ChP, NC = neural crest. (D) Dot plot showing average expression and percentage of cells for key marker genes present in the subclusters identified by scRNA-seq. Lipid-producing ChP express SARS-CoV-2 entry genes ACE2, TMPRSS2 and TMPRSS4. (E) Feature plots showing expression of ACE2 and TMPRSS2 in the subpopulations identified by scRNA-seq. (F) Immunoblot for ACE2 and the loading control β-actin of ChP and telencephalic organoid protein lysates. (G) Representative confocal images of ChP tissue immunostained for the ChP epithelial marker HTR2C (serotonin receptor 2C) in magenta and ACE2 in green. Nuclei in blue are stained with DAPI. Scale bar: 50 μm. (H) gProfileR (Reimand et al., 2011) analysis of genes co-expressed with ACE2 showing significant enrichment (p<0.05) for GO categories cellular component (GO:CC), molecular function (GO:MF) and biological process (GO:BP).

To better characterise the distribution of SARS-CoV-2 entry factors in the different ChP populations, we performed sub-clustering of all the ChP cell populations and we identified four prominent clusters: immature ChP, mature ChP, hem and ChP stroma, as well as two smaller clusters, one that appeared to be neural crest cells, and one that was enriched in expression of lipoprotein genes, suggesting a function in lipid-production and transfer (Fig. 1C). We then looked at the main SARS-CoV-2 entry genes in these cell populations and found expression of entry factors ACE2, TMPRSS2 and TMPRSS4 in the lipoprotein-expressing cells in the ChP, which express genes involved in lipid metabolism such as APOA1 and APOA2 (Besler et al., 2012; Wang et al., 2019) (Fig. 1D-E, Fig. S1B). Next, we validated ACE2 expression by immunoblot comparing telencephalic organoids with ChP organoids (Fig. 1F). We could detect ACE2 in ChP organoid lysates and, to a lesser extent in telencephalic organoids, which would also be expected to contain some telencephalic ChP (Fig. 1F). Next, to better compare expression of ACE2 in both neuronal and ChP tissue simultaneously, we performed immunostaining of organoids containing cortical (Fig. S1C) or ChP identities (Fig. 1G). Consistent with the scRNA-seq data, the ChP epithelial tissue, marked by the serotonin receptor HTR2C, showed sparse but strong positive signal for ACE2 compared to cortex. Together, these findings indicate that ACE2 is expressed in cells of the ChP, but no specific expression of ACE2 is present in neuronal progenitors or neurons. Expression of TMPRSS2 was also detected by immunostaining in the membrane of ChP epithelial cells from organoids (Fig. S1D).

To further explore the expression profile of these identified ACE2 expressing ChP cells, we investigated which genes showed correlated expression pattern with ACE2. Gene Ontology Analysis of genes coexpressed with ACE2 revealed enrichment in GO:MF categories “exogenous protein binding”, “viral receptor activity”, “toxic substance binding” and “aminopeptidase activity”; GO:BP category “protein-lipid complex subunit organization” and GO:CC category “brush border” (Fig. 1H). We found enriched expression of DPP4 (dipeptidyl peptidase 4) and ANPEP (alanyl aminopeptidase), which both encode for known receptors of human CoVs (Qi et al., 2020), and were similarly enriched in the lipoprotein producing subcluster, but not in neurons or other cell types (Fig. S1E, Fig. S1F). In particular, DPP4 is the receptor for MERS-CoV, whereas ANPEP is a receptor for human coronavirus 229E, among other human CoVs (Qi et al., 2020). Interestingly, ChP cells of this subcluster also express lipoproteins that are also required for assembly of hepatitis C virus (HCV), as well as some HCV entry factors such as SCARB1 and CD81 (Aizawa et al., 2015; Grassi et al., 2016; Wrensch et al., 2018) (Fig. S1G). Together these data suggest that: 1. ChP cells and, specifically, lipoprotein-expressing ChP cells, express entry factors required for SARS-CoV-2 infection and 2. Neuronal progenitors and neurons do not specifically express SARS-CoV-2 entry factors.

### SARS-CoV-2 spike pseudovirus only infects ChP cells of brain organoids

To examine SARS-CoV-2 neurotropism, we incubated brain organoids with mixed identities including ChP and cortical tissue with SARS-CoV-2 spike pseudovirions carrying a 19 amino acid deletion (c19) from the C-terminus, which allows for better expression and integration of the spike into the lentivirus (Giroglou et al., 2004; Ou et al., 2020). Spike pseudovirions allow investigation of viral entry without other effects of live CoV such as cell death or viral replication, and because they encoded for GFP we could visualise infected cells only (Giroglou et al., 2004). As a negative control, pseudovirions lacking viral envelope glycoprotein (Δenv) were used for the infection. VSV-G pseudotyped lentivirus (VSV) with broad viral tropism was used as a positive control (Fig. S2A-C). A biochemical assay of the viral reverse transcriptase (RT) was used to assess viral particle production and subsequent titration in ACE2 overexpressing 293T cells (Fig. S2A-C). We initially examined infection by direct observation of GFP fluorescence (Fig. S2D), which clearly showed positive cells, but also some putative autofluorescence in the green channel. Indeed, we found that dead or dying cells displayed quite bright fluorescence in the same channel as GFP (Fig. S2E). Therefore, to be sure we were observing true GFP signal from infected cells, we added a step of immunostaining with a GFP antibody, which allowed for accurate assignment of infected cells.

We performed three independent infections with SARS-CoV-2 spike pseudovirions of organoids, which revealed viral tropism for ChP epithelial tissue, as indicated by the GFP antibody-positive signal (Fig. 2A). Positive control for the infection with VSV lentivirus confirmed a broader tropism (Fig. 2B, Fig. S2F), whereas no positive signal was detectable in organoids infected with Δenv lentivirus (Fig. 2C, Fig. S2F). Quantification of the ratio of GFP-positive cells to total cells showed that spike pseudovirions infected around 13% of cells in the ChP epithelium (Fig. 2D).

**Figure 2.**
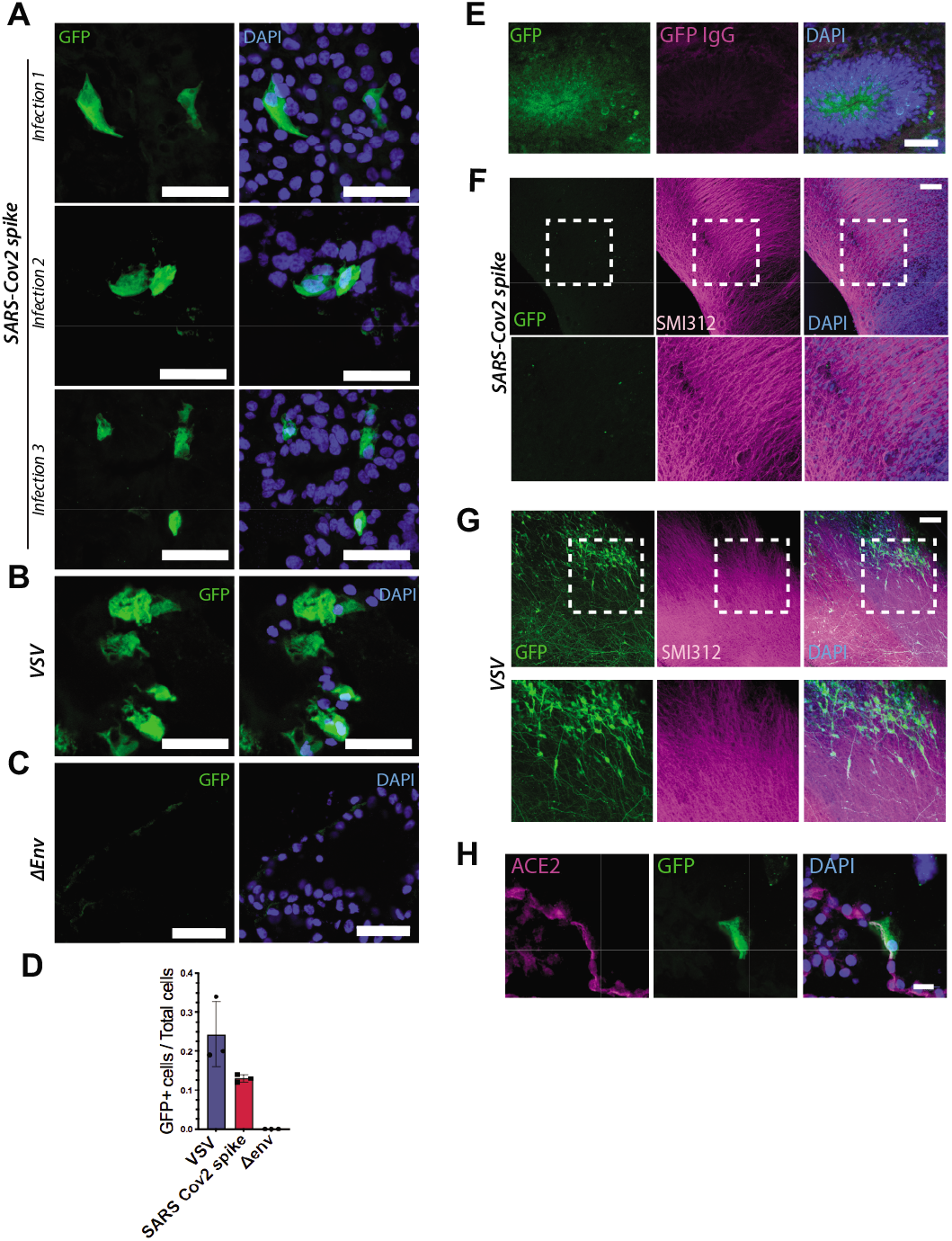
SARS-CoV-2 spike pseudovirus infects ChP cells but not other brain cells of cerebral organoids. (A) Representative confocal images of ChP epithelial tissue from organoids infected SARS-CoV-2 spike pseudovirions. GFP positive cells, as detected by GFP antibody, are shown from three independent experiments. Nuclei in blue are stained with DAPI. Scale bar: 50 μm. (B) Representative confocal images of ChP epithelial tissue from organoids infected with VSV-G lentivirus. Scale bar: 50 μm. (C) Representative confocal images of ChP epithelial tissue from organoids infected with lentivirus pseudovirions lacking viral glycoprotein of the envelope (Δenv). Scale bar: 50 μm. (D) Quantification of GFP-positive cells over total counted cells for VSV-G, SARS-CoV-2 spike and Δenv lentiviral infected ChP epithelial cells from organoids. (n=100 cells counted for each of the three independent experimental repeats). (E) Representative confocal image of a cortical lobe of a cerebral organoid infected with spike pseudovirions showing an example of a false positive signal due to GFP autofluorescence and stained with anti-GFP antibody (in magenta). Nuclei in blue are stained with DAPI. Scale bar: 50 μm. (F) Representative images of ALI-CO infected with spike pseudovirions and immunostained with axonal marker SMI312 in magenta, anti-GFP antibody in green, and DAPI in blue. Scale bar: 100μm. (G) Representative images of ALI-CO infected with VSV lentivirus and immunostained with axonal marker SMI312 in magenta, anti-GFP antibody in green, and DAPI in blue. Scale bar: 100 μm. (H) Higher magnification image of ChP epithelial tissue from organoids immunostained for ACE2 in magenta, GFP and DAPI. Scale bar: 20 μm.

Interestingly, we found that neuronal regions of organoids with mixed identity did not seem to get infected with the virus (Fig. S2G), and the only signal we could detect in the green channel was autofluorescence (Fig. 2E, Fig. S2G). To further investigate the ability of spike pseudovirus to infect cortical tissue and neurons, we infected Air-Liquid Interphase Cerebral Organoids (ALI-COs) (Giandomenico et al., 2019), which are long-term cultures of organoids that lead to improved neuronal maturity and function. ALI-COs infected with spike pseudoviruses and immunostained for SMI312 to visualise the neurons showed no specific GFP-positive signal (Fig. 2F), when compared to the signal detected in ALI-COs infected with the positive control (Fig. 2G). This indicated the lack of signal upon SARS-CoV-2 spike infection was not due to defective GFP expression, but rather a lack of viral entry.

These findings support the hypothesis that susceptibility to infection by SARS-CoV-2 in the brain is also governed by expression of ACE2, which is only present on ChP epithelial cells of the organoids. Further supporting this claim, confocal imaging of infected cells immunostained for ACE2 showed positive signal on the membrane of spike pseudoviruses infected ChP cells (Fig. 2H).

### Live SARS-CoV-2 productively infects ChP tissue

Recent reports have similarly explored neurotropism of SARS-CoV-2 using brain organoids and neurospheres, but with contradictory findings to those we observed here (Ramani et al., 2020; Song et al., 2020). One possible explanation is the use of pseudovirus versus live SARS-CoV-2, with other reports making use of live viral isolates. Therefore, to explore this possibility we turned to infections with a live SARS-CoV-2 clinical isolate (Papa et al., 2020). We first infected ChP organoids with an equivalent amount of virus as we had performed using pseudovirions, and observed highly comparable infection (Fig. 3A, B). We next tested whether this same amount of virus was capable of infecting neurons and other cortical cells of telencephalic organoids with a mixed identity containing also ChP. We still found infection exclusively of the ChP in these conditions (Fig. 3C). Again, this infection matched ACE2 expression, with infected cells co-staining for the receptor (Fig. 3D) in the ChP epithelium, whereas stromal cells of the ChP were uninfected (Fig. S3).

**Figure 3.**
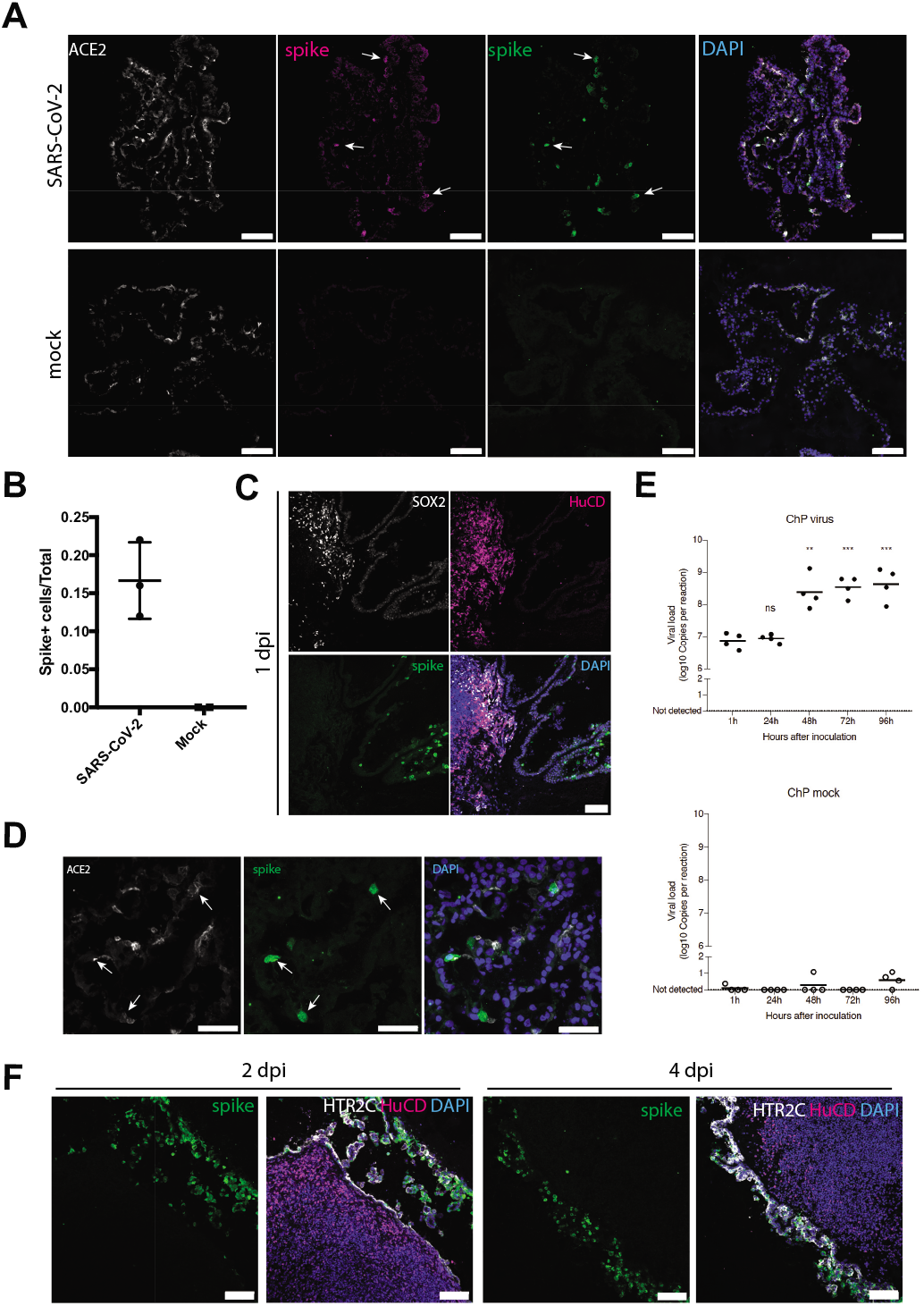
Live SARS-CoV-2 productively infects ChP epithelium. (A) 1 day post-infection of ChP organoids with either live SARS-CoV-2 or mock and staining for two independent antibodies (Abcam ab252690 spike glycoprotein in magenta and GeneTex GTX632604 in green) directed to the viral spike protein. Specific staining (arrows) is only seen in the SARS-CoV-2 infection condition. Scale bars: 100 μm. (B) Quantification of infected cells staining positive for viral spike protein in ChP tissue infected with SARS-CoV-2 (n=3 independent infections) compared with mock (n=2 independent infections). (C) Staining for viral spike protein in mixed identity telencephalic organoids at 1 day post-infection (dpi) showing staining only in ChP tissue. Scale bar: 100 μm. (D) Co-staining for ACE2 in infected cells of ChP tissue (arrows). Scale bars: 50 μm. (E) RT-qPCR using primers and probes against the CDC N1 amplicon of SARS-CoV-2 in infected and mock over the course of 4 days post-infection. n=4 organoids from 2 independent infections. \*\**P*=0.002, \*\*\**P*<0.001, Two-tailed unpaired Student’s *t*-test. (F) Staining for viral spike protein in mixed identity telencephalic organoids displaying adjacent cortical and ChP tissues at 2 days post-infection and 4 days post-infection. Scale bars: 100 μm.

We next examined whether ChP cells represent a permissive cell type for viral replication. We performed qRT-PCR for viral N1 and observed a significant increase in viral genome copies in organoid supernatant over the course of 4 days (Fig. 3E) with the largest increase between 1 and 2 days of infection. This matched staining where we similarly observed larger numbers of cells infected in the ChP at 2 and 4 days post-infection (Fig. 3F).

### SARS-CoV-2 has minimal tropism for neurons but disrupts barrier integrity of the ChP

We next sought to test whether there could be any conditions in which neurons or other CNS cell types may be infected with SARS-CoV-2. We did not observe any viral spreading from ChP to nearby cortical regions, even when in very close proximity and left for as long as 4 days (Fig. 3F). However, one possibility is that the virus preferentially infects ChP resulting in lower amounts of viral particles available to infect the surrounding neurons. We therefore tested infection with live virus on “pure” cortical organoids containing no ChP, but still saw no specific staining for viral protein in neurons or neural progenitors (Fig. 4A, Fig. S4A). Finally, we tested whether a combination of longer exposure and much higher viral quantities would lead to infection. We performed infection of ALI-COs with 10 times the viral MOI used for ChP infection, and were able to observe very sparse, but specific, neuronal and glial infection after 2 days post-infection (Fig. 4B, Fig. S4B). This suggests that SARS-CoV-2 has a much lower infectivity of human neural cell types compared with ChP epithelium. This finding was also consistent with the expression of markers of the lipoprotein-expressing cell type, which were present in cells infected by SARS-CoV-2 in the ChP epithelium (Fig. 4C, D), but absent from cortical neuronal regions (Fig. 4E).

**Figure 4.**
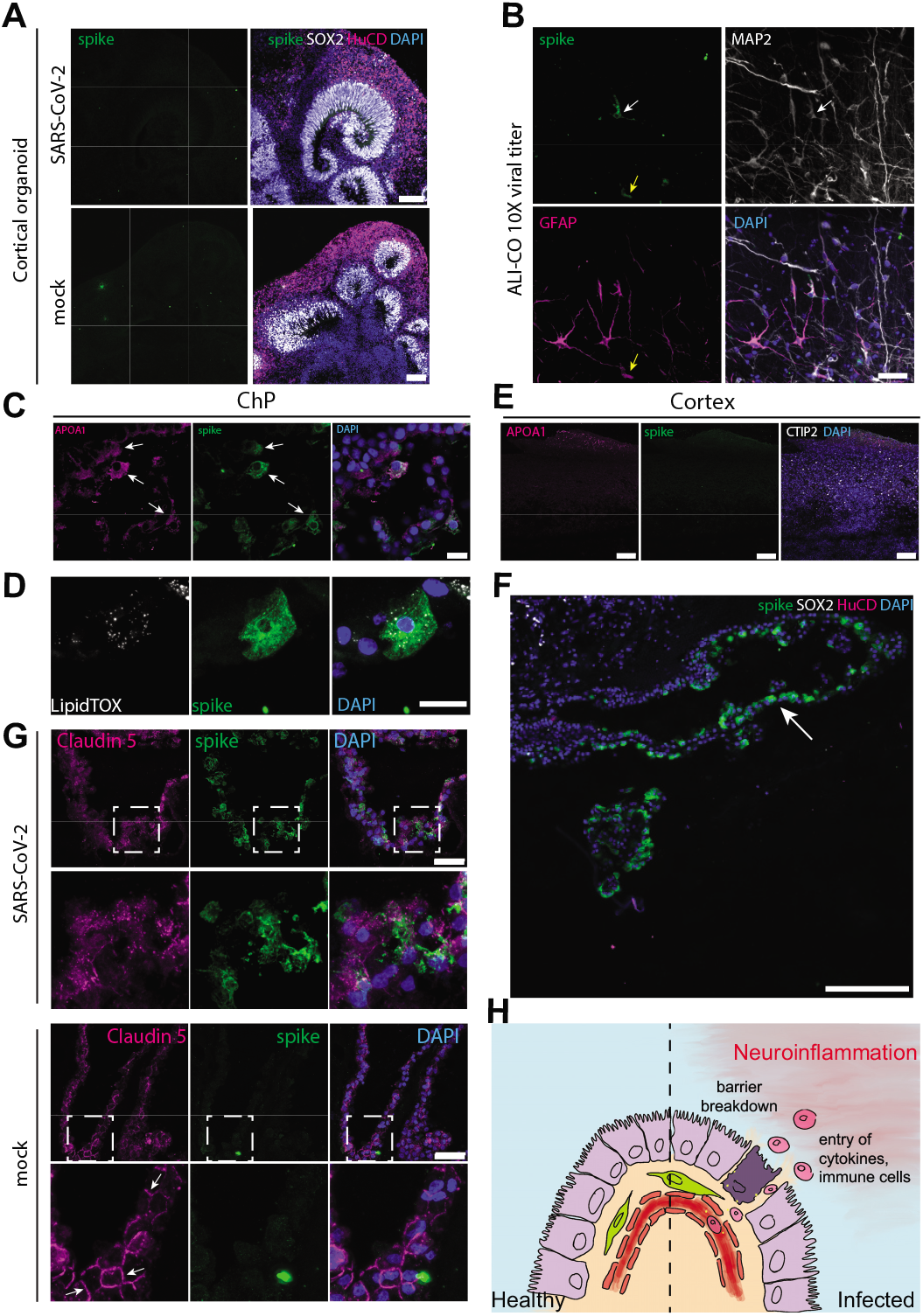
SARS-CoV-2 disrupts ChP epithelial integrity. (A) Staining for viral spike protein in pure cortical organoids infected with either SARS-CoV-2 or mock showing no specific staining in either condition. Scale bars: 100 μm. (B) Staining for viral spike protein in ALI-COs infected for 2 days with 10 times the dose used for ChP or mixed identity organoids. Note sparse staining of a neuron (white arrow) that is positive for MAP2 and a glial cell (yellow arrow) that is positive for GFAP. Scale bars: 50 μm. (C) Co-staining for APOA1 and viral spike protein in ChP epithelium after 1 day post-infection. Scale bar: 20 μm. (D) Co-staining for LipidTOX and viral spike in a ChP epithelial cell after 1 day post-infection. Scale bars: 20 μm (E) APOA1 staining in infected cortical tissue showing no viral spike staining and only sparse APOA1 outside the cortical neuronal region marked by CTIP2. Scale bars: 100 μm. (F) Staining for viral spike protein in telencephalic organoid with intact CSF-like fluid filled cyst (arrow) showing infection after application of virus on the basal (outer) surface. Scale bar: 200 μm. (G) Staining for tight-junction protein Claudin 5 in ChP epithelium infected with SARS-CoV-2 or mock at 2 days post-infection. Note the presence of clearly demarcated junctions (arrows) in mock versus infection with SARS-CoV-2. Green fluorescent signal in mock represents typical nonspecific background, which does not label a cell. Scale bars: 50 μm. (H) Model of the effect of SARS-CoV-2 infection of the ChP and the effect on CNS barrier integrity.

We next examined the impact of SARS-CoV-2 infection on the ChP epithelium. First, because the ChP epithelium is highly polarized with its basal side facing the endothelial compartment, potential viral entry in vivo would have to occur from the basal surface. We therefore tested whether SARS-CoV-2 could successfully infect the ChP epithelium from the basal, or media side, of intact tissues with clear, fluid-filled cysts. This experiment revealed abundant infection from the basal side (Fig. 4F), suggesting that blood-borne virus can indeed infect from the vascular compartment of the ChP. Second, because the ChP is an integral part of the blood-CNS barrier, we examined the effect of viral infection on barrier integrity. We observed a striking effect on cell-cell junctions (as labeled by Claudin-5) after 2 days post-infection (Fig. 4G) and after 4 days the tissue was visibly damaged compared with mock (data not shown). Taken together, these data suggest that SARS-CoV-2 is capable of infecting a subset of ChP epithelial cells, which leads to a breakdown of barrier integrity (Fig. 4H).

## Discussion

Preliminary clinical reports on SARS-CoV-2 have highlighted neurological symptoms such as headache, encephalitis and nausea, suggesting a neuroinvasive potential of the virus (Montalvan et al., 2020; Moriguchi et al., 2020). It was unclear, however, whether SARS-CoV-2 can infect neuronal cells. Understanding viral tropism is of significant interest for the development of better treatments and for prevention of long-term adverse effects. Our findings indicate that SARS-CoV-2 does not readily infect neuronal cells, but rather infects ChP epithelial cells of the brain. This finding is consistent with the high levels of expression of SARS-CoV-2 entry factors such as ACE2 and TMPRSS2 in the ChP *in vivo* and *in vitro*, compared to other brain regions (Chen et al., 2020). Furthermore, our findings of susceptibility of a specific lipoprotein-producing cell type, matches closely the susceptibility seen in other organs such as the intestine (Lamers et al., 2020).

A likely interpretation of these results is that the neurological symptoms reported in COVID-19 patients are mainly due to an indirect, secondary consequence of viral infection of support cells in the brain, rather than neurons themselves. We show that infection of the ChP epithelium by SARS-CoV-2 leads to disruption of the B-CSF-B. While this could then allow entry and spread of the virus into the brain, our results suggest that neural cells are minimally susceptible, even when exposed to high quantities of virus. Furthermore, substantial SARS-CoV-2 within the brain and CSF does not seem to be a widely reported finding (Schaller et al., 2020). Nonetheless, barrier breakdown could allow abnormal entry of immune cells and cytokines leading to harmful neuroinflammation (**Fig. 4H**).

It is known that immune cells, including macrophages, dendritic cells and monocytes, continuously survey the B-CSF-B, and infection of ChP epithelial cells can lead to pro-inflammatory cytokine production and recruitment of T-cells that can initiate neural tissue injury (Ransohoff and Engelhardt, 2012). Furthermore, the B-CSF-B might be a particularly vulnerable barrier to invasion of this sort because the tight junctions between cells have lower electrical resistance compared to the blood-brain-barrier (BBB) (Redzic, 2011; Schwerk et al., 2015). Finally, normal functioning of the ChP is necessary for proper CSF production and turnover, which is essential in providing nutrients and clearing metabolic waste. These roles have recently gained attention with the discovery of the so-called “glymphatic system”, which seems to be most active during sleep when changes in CSF flow also occur (Fultz et al., 2019; Olsson et al., 2018).

It is interesting to note that among the neurological disturbances experienced in this and previous CoV outbreaks, chronic fatigue and nonrestorative sleep disturbances have been noted (Moldofsky and Patcai, 2011). Indeed, multiple viruses, such as HCV, HIV and EBV, have been linked to chronic fatigue syndrome (CFS), also known as myalgic encephalomyelitis (ME) (Payne et al., 2013; Rasa et al., 2018; Williams et al., 2019). In the case of HCV, the most common extrahepatic symptom in patients is chronic fatigue (Poynard et al., 2002). Key mediators of HCV assembly are apolipoproteins, and regulation of lipoprotein metabolism is important for the HCV replication cycle (Aizawa et al., 2015; Grassi et al., 2016). Lipoproteins are mainly synthesized by hepatocytes and intestinal cells (Aizawa et al., 2015); however, we report here a previously unrecognized lipoprotein-producing cell population in the ChP that also expresses viral entry factors for HCV and for SARS-CoV-2, perhaps explaining the particular susceptibility of this CNS tissue.

While the exact cause of CFS/ME is still obscure, abnormal levels of a number of cytokines, such as IL-10 have been observed in CSF from patients with CFS/ME (Natelson et al., 2005), suggesting an imbalance in neuroimmune modulation. This points to a potential breakdown or dysfunction of the B-CSF-B, allowing leakage of immune cells and cytokines into the brain. Alternatively, or perhaps in combination, the role of the ChP in CSF production and turnover could be compromised, thus leading to defects in waste clearance during sleep.

Together, these observations, combined with our results from human brain organoids, suggest the intriguing possibility that the ChP epithelium is particularly susceptible to infection by viruses, and that this susceptibility may lead to breakdown of the B-CSF-B and development of CFS/ME symptoms. In the case of SARS-CoV-2, this susceptibility seems to be dictated by the expression of ACE2, which is primarily expressed in ChP epithelium. Thus, our findings suggest that the neurological symptoms are not due to a direct effect on neurons, but rather a consequence to the damage of this important supportive tissue resulting in proinflammatory changes in the fluid it secretes.

## Materials and Methods

### Cells and plasmids

HEK 293T CRL-3216 cells were purchased from ATCC and authenticated by the supplier. All cells were regularly tested and were mycoplasma free. HEK 293T cells were maintained in Dulbecco’s modified Eagle’s medium (DMEM) with 10% FBS, 2 mM Lglutamine, 100 U/ml penicillin, and 100 μg/ml streptomycin (Gibco) at 37°C with 5% CO_2_. Vero cells were purchased from ATCC and maintained in DMEM supplemented with 10% FBS, 100 U/ml penicillin and 100mg/ml streptomycin. Cells tested negative for mycoplasma before virus production and infection experiments. Vero-hACE2 were generated by transducing Vero cells with lentiviral particles expressing hACE2 ORF and cultured in DMEM 10% FCS with 5 μg/ml blasticidin. Vero-hACE2-TMPRSS2 were generated by transducing Vero-hACE2 with lentiviral particles expressing TMPRSS2 ORF and maintained in DMEM 10% FCS with addition of 5 μg/ml blasticidin and 1 μg/ml puromycin. H9 ES cells were obtained from WiCell (WA09) and have been approved for these studies by the UKSCB Steering Committee. Human ES cells were maintained in culture with StemFlex culture media (Gibco A3349401) on growth factor reduced Matrigel coated dishes (Corning).

Vectors for viral production were: pCRV-1 for HIV-1 Gag/Pol (Naldini et al., 1996), and CSGW for GFP expression (Zennou et al., 2004). Lentiviral packaging plasmid pMDG2, which encodes VSV-G envelope, was used to pseudotype infectious virions (Addgene plasmid # 12259). pCAGGS-Spike Δc19 was generated from pCAGGS-Spike by digesting the vector backbone with EcoR1 and NheI and subsequently gel purified. Q5 polymerase (NEB) was used to amplify the spike protein fragment using the 3’ primer to make a C-terminal 19 amino acid deletion. Primers with a 15bp overlap with the vector backbone were used. After PCR, the fragments were treated with Dpn1 for 1h and subsequently gel purified. The fragment was then inserted into the vector using In-Fusion assembly (Takara Inc.). The plasmid was checked by sequencing.

For the generation of ACE2 over-expressing HEK 293T cells, the human ACE2 ORF was PCR amplified from Addgene plasmid 1786 and C-terminally fused with the porcine teschovirus-1-derived P2A cleavage sequence (ATNFSLLKQAGDVEENPGP) followed by the blasticidin resistance gene. This continuous, single ORF expression cassette was transferred into pLenti6-Dest_Puro by gateway recombination. Lenti-viral particles were generated by cotransfection of HEK 293T cells with pLenti6-Dest_Puro_ACE2-2A-Bla, pCMVR8.74 (Addgene plasmid 22036) and pMD2.G (Addgene plasmid 12259) using PEI. Supernatant containing virus particles was harvested after 48 h, 0.45 um filtered, and used to infect naive HEK 293T cells. Transduced cells stably expressing ACE2 were selected with 5 ug/ml blasticidin.

### Preparation of viruses

Replication deficient VSV-G or SARS Cov2 pseudotyped HIV-1 virions were produced in HEK293T cells by transfection with pMDG2 or pCAGGS-Spike Δc19, pCRV GagPol and CSGW as described previously (Price et al., 2014). Viral supernatants were filtered through a 0.45μm membrane at 48 hours post-transfection and pelleted through a 20% sucrose cushion for 2hrs at 28K. Pelleted virions were drained and then resuspended in DMEM. Viral stocks were quantified by qRT-PCR as described previously (Vermeire and Verhasselt, 2013) with modifications. In brief, 5 μl of virus stock was mixed with 5 μl lysis buffer (0.25% Triton X-100, 50 mM KCl, 100 mM Tris-HCl (pH 7.4), 40% glycerol, 0.1 μl RNase Inhibitor). After incubation at room temp for 10 mins, 90 μl nuclease-free water was added. 2 μl of lysate was mixed with 5 μl TaqMan Fast Universal PCR Mix, 0.1 μl MS2 RNA, 0.05 μl RNase Inhibitor and 0.5μl MS2 primer/probe mix (7.5μM primers Fwd: AACATGCTCGAGGGCCTT, Rev: GCCTTAGCAGTGCCCTGTCT, 3.7μM probe, TGGGATGCTCCTACATG), in a final volume of 10μl.

For titer determination of pseudotyped viruses, 293T ACE2 cells were plated into 96 well plates at a density of 7.5×10^3^ cells per well and allowed to attach overnight. Viral stocks were titrated in triplicate by addition of virus onto cells. Infection was measured through GFP expression measured by visualisation on an Incucyte Live cell imaging system (Sartorius). Infection was enumerated as GFP positive cell area.

Live SARS-CoV-2 (SARS-CoV-2/human/Liverpool/REMRQ0001/2020) used in this study was isolated by Lance Turtle (University of Liverpool), David Matthews and Andrew Davidson (University of Bristol). SARS-CoV-2 stock was prepared in Vero hACE2-TMPRSS2 cells by infecting monolayer of cells with 0.01 MOI of virus. Virus stock was harvested after 3 days by three freeze-thaw cycles and 5 min 300xg spin to remove cell debris. Virus titers were assessed by plaque assays in Vero ACE2/TMPRSS2 cells. For plaque assays, Vero hAce2-TMPRSS2 cells were seeded on 12-well dishes day prior infection. Next day serial dilutions of the supernatant (−1 to −6) were prepared and used to infect cells for 1h and then overlayed with 0.05% agarose in 2% FBS DMEM. After 3 days monolayers were fixed with 4% formaldehyde and the plaques were revealed with 0.1% toluidine blue staining.

### Cerebral and ChP organoid culture condition

Cerebral organoids were prepared from single cell suspension of human ES cells as previously described (Lancaster and Knoblich, 2014) and cultured with Stem Cell Technologies Cerebral Organoid kit (catalogue n. 08570, 08571). Briefly, EBs were generated by seeding 2000 cells in a 96-well U-bottom low attachment plate with EB media and 50μM Y-27632 ROCK inhibitor for 3 days. EB media was replaced on day 5 with NI media in the same 96 well plate. At day 7, EBs were embedded in 30μl Matrigel (Corning) using sheets of dimpled parafilm and incubated for 20min at 37°C as previously detailed (Lancaster and Knoblich, 2014). EBs were then transferred to a 6-well plate in 3ml of Expansion media per well. From day 30, dissolved Matrigel (1:50) was added to the Maturation media.

Air-Liquid Interface Cerebral Organoid (ALI-CO) culture was performed as previously described (Giandomenico et al., 2019). Briefly, mature organoids (day 55) were collected, washed in HBSS without Ca2+ and Mg2+ and embedded in 3% low-gelling-temperature agarose (Sigma, A9414) at 40°C in embedding molds (Sigma, E6032). Agarose blocks were then incubated for 10-15min on ice and sectioned using a vibratome (Leica VT100S vibrating microtome) in cold HBSS. Sections (300μm-thick) were collected onto Millicell CM culture inserts (Millipore, PICM0RG50) in 6-well plates and left at 37°C for 1h to equilibrate with serum-supplemented slice culture media (SSSCM). SSSCM was then replaced with serum-free slice culture media (SFSCM). ALI-CO cultures were fed with SFSCM daily.

### Single-cell RNA-seq and in vivo expression analyses

scRNA-seq data was previously published (Pellegrini et al., 2020) and was analysed using the Seurat v3 R package. The already normalized and scaled matrix was visualized by UMAP dimensionality reduction. Subclustering of ChP cell types was performed by first extracting and combining cells of the ChP mature epithelial, immature/hem, and stromal clusters, followed by FindNeighbors, FindClusters and UMAP dimensionality reduction on PCs 1-12 (as determined by ElbowPlot). For expression correlation analysis, pearson correlation was performed between ACE2 expression in all ChP cells and all other genes. The top 20 correlated genes were used for GO term enrichment analysis using gProfiler (Reimand et al., 2011).

*In vivo* data were obtained from the Allen Human Brain Atlas (http://human.brain-map.org/). Z-scaled data were downloaded for the two ACE2 probes available for all samples and brain regions. Mean Z-score was calculated for all samples within each brain region for each probe.

### Organoid infection experiments

Organoid infections were performed by addition of virus to the culture medium at an expected MOI of 0.5 for each virus based on viral titre determined in 293T ACE2 cells (in the case of pseudotyped virus) or plaque assay (in the case of live SARS-CoV-2). To calculate an MOI, counts from previous single cell dissociations (Pellegrini et al., 2020) were used to estimate cell numbers in organoids. Thus, 7×10^3^ PFU virus was added to choroid plexus tissues with an estimated number of 17,000 cells to achieve MOI 0.5. For mixed identity or pure cortical organoids, with an increased cell number estimated at approximately 35,000 cells, we used 1.4×10^4^ PFU virus, or 1.4×10^5^ in experiments using 10-fold increased viral quantity. In the case of organoid ChP tissue, ChP epithelium was physically separated from the organoid and broken into three large pieces before infection, and SARS-CoV-2 virus or pseudovirions, negative control, or positive control was added to each of the three pieces. This provided an added internal control so that the experiment and control conditions were tested on tissues obtained from the exact same organoid.

### qRT-PCR analysis of viral replication

RNA extractions from 140μl of inoculated culture medium were performed using the Qiagen QIAmp Viral RNA mini kit and eluted in 80μl of AVE elution buffer. 5μl of extract were used per 20μl RT-qPCR reaction. Reaction mixes were set up according to the manufacturer’s instructions with the addition of a CDC-N1 SARS-CoV-2 probe/primer mix (IDT 2019-nCoV RUO kit) with a final concentration of 500nM for each primer and 125nM of the hydrolysis probe. One-step RT-qPCR reactions were run on a Viia7s real-time qPCR instrument with the following parameters: 1. 55°C for 10 minutes for reverse transcription 2. 95°C for 1 minute for enzyme activation. 3. 45 cycles of 10 seconds denaturation at 95°C and 30 seconds annealing/extension at 55°C. Fluorescence was recorded at the end of the extension step. To estimate viral copies per reaction, a dilution series of nCoV-2019 N-gene positive control (IDT) was performed in technical duplicates. Sample Cq values were then converted to viral copies per reaction using the standard curve’s linear regression model. Values in dot plots represent log10 transformed copies per reaction of individual biological samples over time. For samples where no virus was detected, a pseudo value of 0 was assigned to represent the data point on the graph. Raw data is available upon request.

### Immunostaining and confocal imaging

Organoids were fixed in 4% PFA overnight at 4°C and then moved to 30% sucrose buffer at 4°C for at least 24h. Organoids were then embedded in gelatin and sectioned as previously described (Lancaster and Knoblich, 2014). After blocking and permeabilisation with 0.25% Triton and 1% donkey serum buffer, sections were incubated overnight with the following primary antibodies according to their optimized instruction: sheep anti-TTR (1:500, Abcam, ab9015), rabbit anti-ACE2 (1:100, Abcam, ab108209), goat anti-ACE2 (1:100, R&D Systems, AF933), rabbit anti-HTR2C (1:500, Abcam, ab133570), mouse anti-TMPRSS2 (with antigen retrieval, 1:50, SantaCruz, Sc515727), rabbit anti-claudin 5 (1:200, ThermoFisher, 34-1600), rabbit anti-Sox2 (1:300, Abcam, ab97959), rabbit anti-APOA1 (1:100, ThermoFisher, PA5-88109), sheep anti-Tbr2 (1:200, R&D Systems, AF6166), rat anti-CTIP2 (1:300, Abcam, ab18465), mouse anti-HuC/D (1:500, LifeTech, A21271), rabbit anti-DLK1 (1:200, Abcam, ab21682), rabbit anti-GFAP antibody (1:200, Abcam, ab7260), chicken anti-MAP2 (1:1000, Abcam, ab5392),_chicken anti-GFP (1:500, Abcam, ab13970), goat anti-MSX1 (1:200, R&D, AF5045), rabbit anti-ND2 (1:500, Abcam, ab104430), mouse anti-SMI312 (1:500, BioLegend, 837934), mouse anti-SARS-CoV-2 spike glycoprotein (1:200, GeneTex, GTX632604), rabbit anti-SARS-CoV-2 spike glycoprotein C-terminal (1:200, Abcam, ab252690). To mark the nuclei, Dapi (1:1000) was used added with the secondary antibody incubation. Secondary antibodies labelled with Alexa 488, 568 and 647 were applied 1h at room temperature. Incubation with HCS LipidTox (1:1000 in PBS, Thermo Fisher, H34477) was carrier for 20min after secondary antibody to visualize lipid droplets. Slides were then washed in PBS and mounted using Prolong Diamond mounting media. All the staining steps for ALI-COs were carried out in permeabilisation buffer and their duration was extended as previously described (Giandomenico et al., 2019).

Images were acquired using a Zeiss LSM 780 confocal microscope (Carl Zeiss) and prepared using Fiji (NIH). The Fiji Huttner plugin was used for the quantification of GFP-positive cells over total cells: 100 Dapi-stained nuclei were counted in each experiment and number of GFP-positive cells over total cells counted was then recorded.

### Immunoblotting

For immunoblotting, organoids were snap-frozen in liquid nitrogen and homogenized in RIPA buffer with protease inhibitors (Roche) to produce total protein lysate. Protein samples were prepared using NuPAGE LDS Sample Buffer 4x and DTT 1M and heated at 95°C for 15min. Protein samples and ladder were loaded into a polyacrylamide gel and run for 2h at 100mV. Wet transfer in PDVF membrane (Immobilion) was carried out for 3h or overnight at 4°C. Membranes were blocked in 5% milk in PBS-T and incubated overnight at 4°C with primary antibodies (rabbit anti-ACE2 and mouse anti-β-actin). After 3 washes in PBS-T, secondary antibodies Alexafluor conjugated were added for 1h at room temperature and membranes were images using a Li-COR Odyssey CLx Infrared Imaging System.

## Author contributions

L.P. designed and conducted experiments, analysed data, and wrote the manuscript. A.A. performed viral experiments and collected samples. D.L.M. prepared and analysed pseudoviruses. M.J.K. performed RT-qPCR analyses. D.P. generated reagents for cell line generation. A.P.C. generated reagents and cloning constructs. L.C.J. designed and supervised viral work. M.A.L. designed and supervised the project, analysed data, and wrote the manuscript.

## Acknowledgments

The authors would like to thank members of the Lancaster lab for helpful feedback and discussions. We also thank G. Papa for cell lines used in these studies. We also thank the Light Microscopy and BSL3 facilities of the MRC Laboratory of Molecular Biology. This work was supported by the Medical Research Council (MC_UP_1201/9).

## Data and materials availability

Single cell sequencing data were previously published and accessible on NCBI GEO, accession number GSE150903, and through the UCSC cell browser. Raw data is available upon request. H9 (WA09) cells are available through MTA with WiCell.

## Supplementary Figures

**Supplementary Figure 1.**
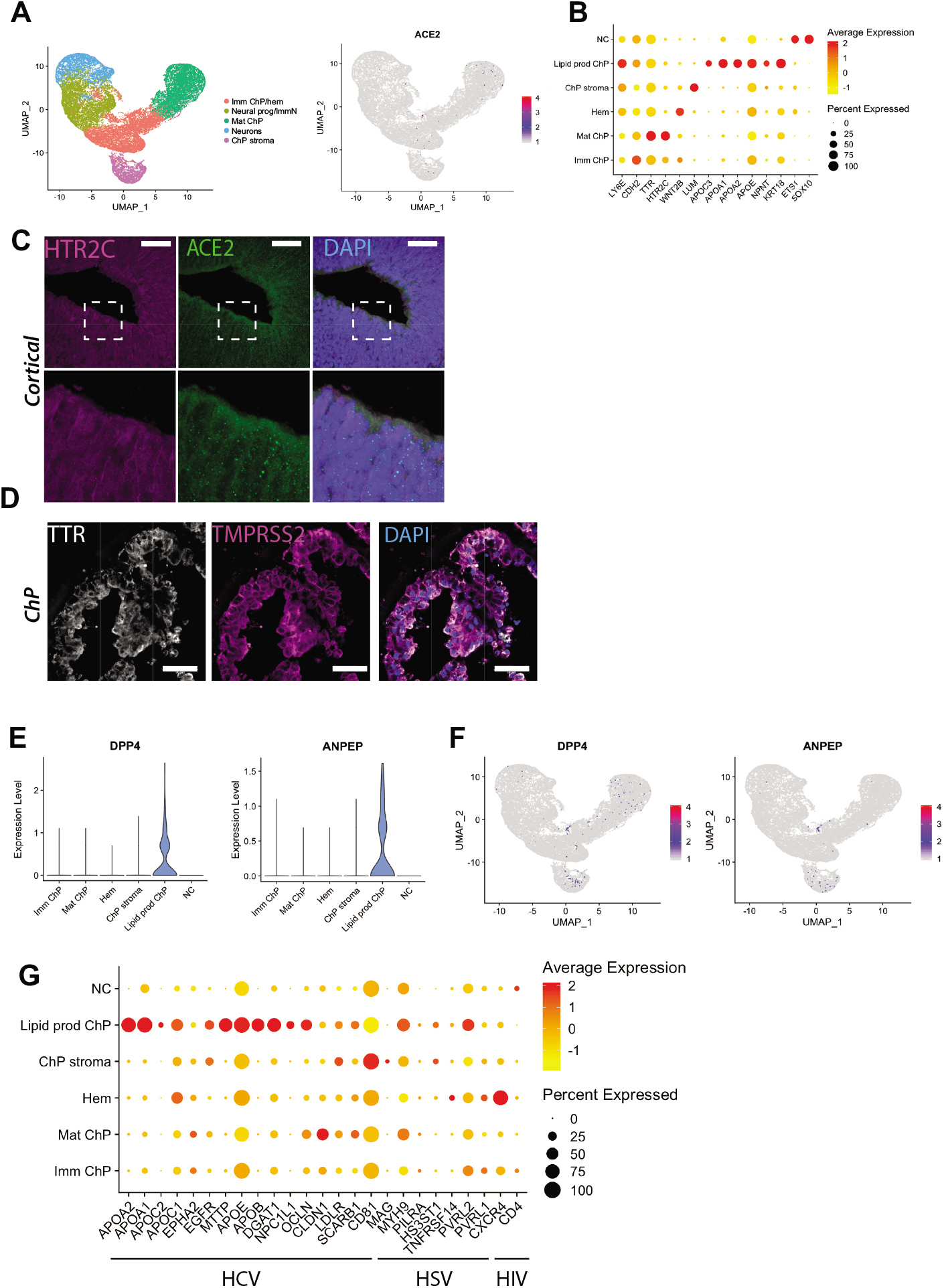
(A) UMAP plot showing the main populations previously identified by scRNA-seq in cerebral and ChP organoids combined (Pellegrini et al., 2020) (left) and feature plot showing sparse expression of ACE2 in the ChP clusters (right). (B) Dot plot showing average expression and percentage of cells expressing top genes in the six main ChP subclusters identified by scRNA-seq. Lipid-producing ChP cells express apolipoprotein genes such as APOC3, APOA1, APOA2 and APOE as well as epithelial markers such as KRT18 (keratin 18) and lateral ventricle ChP marker LY6E (lymphocyte antigen 6E). (C) Representative confocal images of cortical lobe from an organoid with mixed identity immunostained for ChP marker HTR2C (magenta), ACE2 (green) and DAPI (blue). Scale bar: 50 μm. (D) Representative confocal images of a ChP epithelial region from a mixed identity organoid stained for ChP marker TTR (grey), TMPRSS2 (magenta) and DAPI (blue). Scale bar: 50 μm. (E) Violin plots showing expression levels of human coronavirus entry factors DPP4 and ANPEP in the lipid-producing ChP cell subcluster identified by scRNA-seq. (F) Feature plots showing expression of DPP4 and ANPEP in the ChP subpopulations identified by scRNA-seq analysis. (G) Dot plot showing average expression and percentage of cells expressing HCV viral entry factors in the six subclusters identified by scRNA-seq of ChP organoids. ChP lipid-producing cells show the highest expression of HCV viral entry factors as compared to the other subpopulations.

**Supplementary Figure 2.**
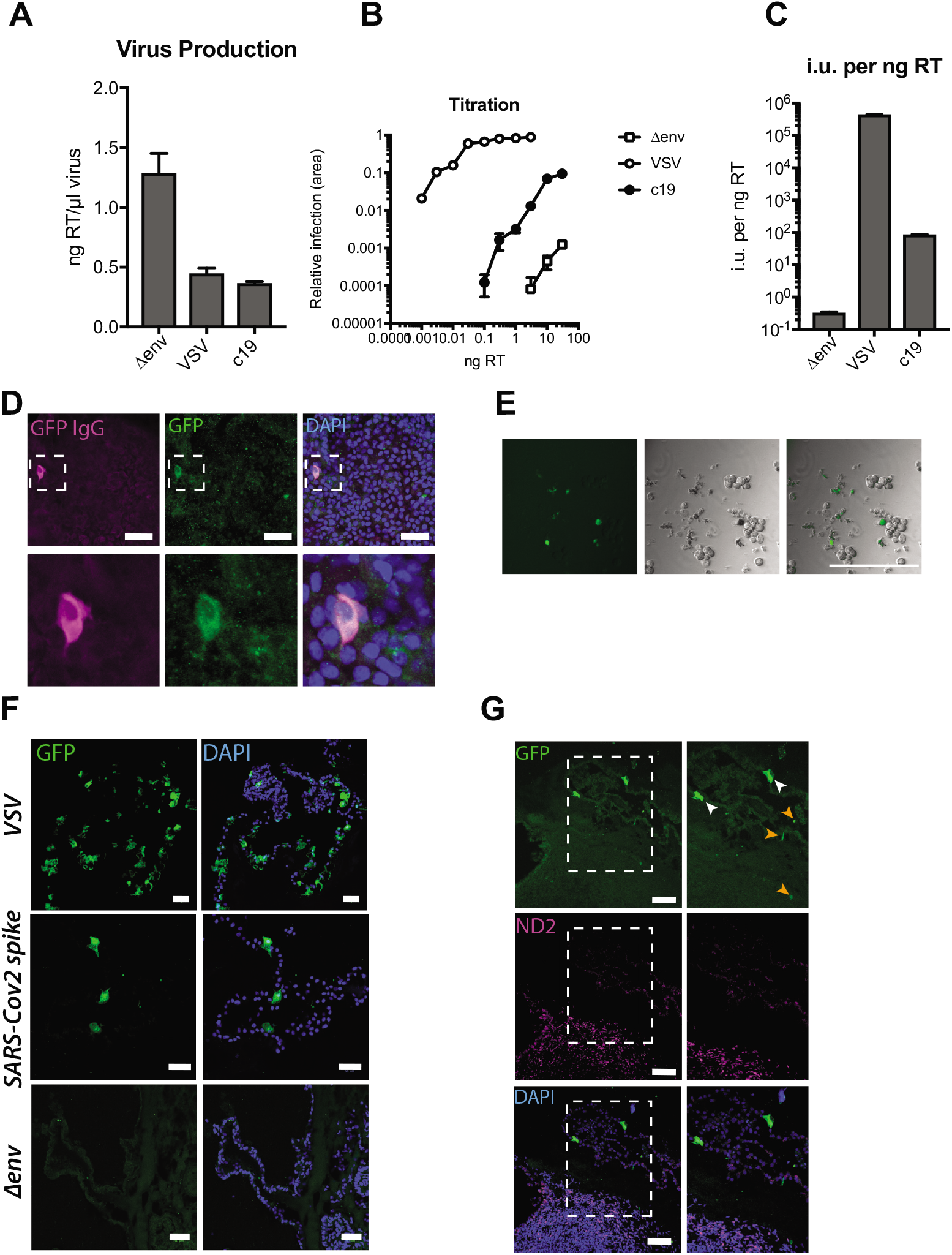
(A) Quantification of viral particle production showing levels of RT enzyme per μl of virus for pseudovirions lacking viral glycoprotein of the envelope (Δenv), VSV lentivirus and SARS-CoV-2 spike with deletion of 19 aminoacids from the C-terminus (c19). (B) Titration of Δenv, VSV and SARS-CoV-2 pseudoviruses onto ACE2 overexpressing 293T cells showing amount of virus added as quantity of RT (ng) on the x-axis. (C) Quantification of infectious units equalised per particle addition (ng RT). (D) Confocal images showing a ChP cell infected with SARS-CoV-2 immunostained with GFP antibody (GFP IgG, in magenta) and DAPI (blue). Scale bar: 50 μm. (E) Epifluorescence images of uninfected, dissociated ChP cells, taken with Evos cell imaging system showing some autofluorescent dead cells. Scale bar: 200 μm. (F) Representative lower magnification confocal images of ChP tissue infected with VSV, SARS-CoV-2 spike and Δenv pseudovirions immunostained for GFP antibody and DAPI in blue. Scale bar: 50μm. (G) Confocal images of an organoid region with cortical and ChP tissue immunostained for neuronal marker ND2 (magenta), GFP and DAPI (blue). White arrowheads indicate GFP-positive cells, orange arrowheads point to autofluorescence possibly derived from cellular debris.

**Supplementary Figure 3.**
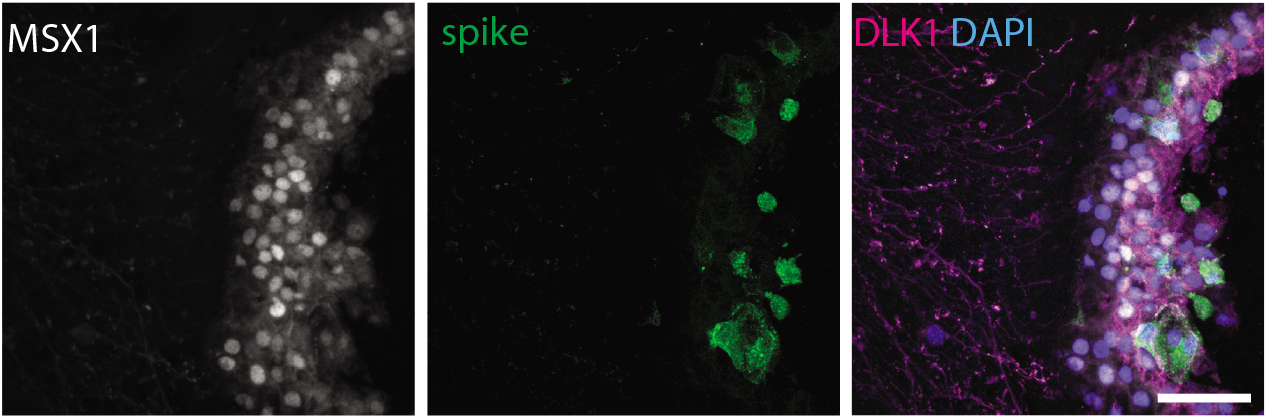
Staining for viral spike protein and a marker of ChP stroma (DLK1) as well as ChP marker MSX1 showing no infection of stromal cells despite abundant infection of epithelium. Scale bar: 50 μm.

**Supplementary Figure 4.**
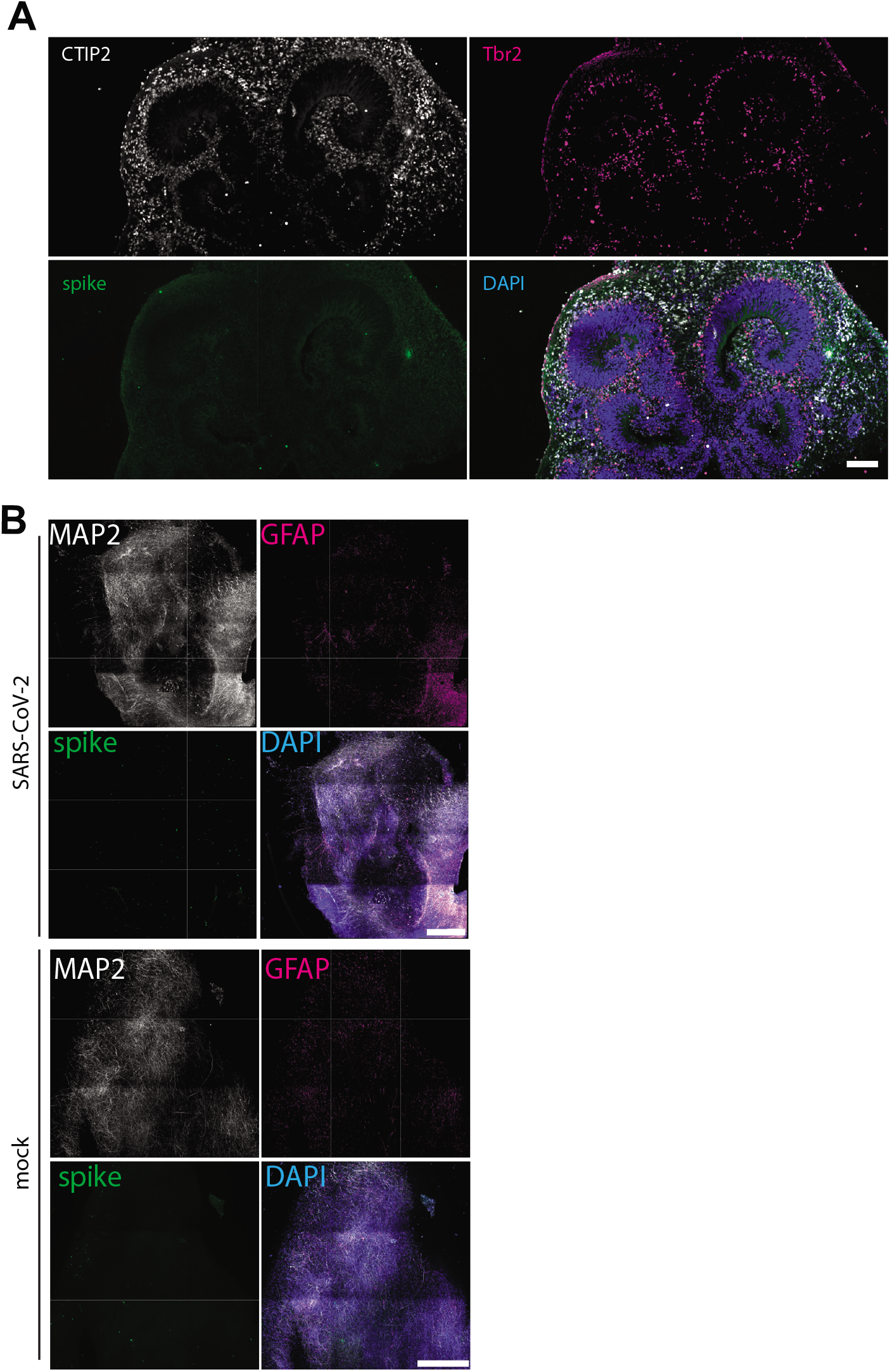
(A) Staining for viral spike protein in a cortical organoid infected with live SARS-CoV-2 at 1 day post-infection showing no specific staining in CTIP2+ neurons or in TBR2+ intermediate progenitors. Scale bars: 100 μm. (B) Overview image of ALI-CO infected with 10 times viral titer at 2 days post-infection showing only very sparse staining, despite abundant neurons (MAP2) and glia (GFAP). Scale bars: 500 μm.

